# Re-evaluating the Impact of Biological Sex on Atherosclerosis in Apoe^−/−^ and Ldlr^−/−^ mice

**DOI:** 10.1101/2025.09.19.677480

**Authors:** Yaping Zhao, Li Wang, Bradford C. Berk, Hans Strijdom, Yu Huang, Jianping Weng, Suowen Xu

**Author notes:** Author for correspondence: Suowen Xu, PhD, Department of Endocrinology, Institute of Endocrine and Metabolic Disease, The First Affiliated Hospital of University of Science and Technology of China, 17 Lujiang Road, Hefei, 230001, China.

## Abstract

**BACKGROUND:** The progression of cardiovascular disease shows significant sexual dimorphism: although females generally develop the disease later in life, they exhibit a higher age-related incidence than males. Despite this clinical pattern, preclinical studies often overlook both sexes in their design, and existing research on sex-specific atherosclerosis in mice remains inconsistent. This study was designed to assess the influence of sex on atherosclerosis using two widely used atherosusceptible mouse models—Apoe^−/−^ and Ldlr^−/−^ mice.

**METHODS:** To investigate the influence of sex on atherogenesis, we conducted a 20-week study using both male and female Apoe^−/−^ and Ldlr^−/−^ mice. The mice were fed an atherogenic ALMN diet (40% trans-fat, 2% cholesterol, 22% fructose) to promote plaque development. We performed comprehensive analyses of: (1) systemic metabolic parameters (lipid profile, glucose metabolism); (2) atherosclerotic burden (*en face* and aortic sinus plaque area); and (3) plaque composition (necrotic core size, collagen content, macrophage infiltration).

**RESULTS:** In both models, male mice showed higher lipid levels, worse glucose tolerance, and reduced insulin sensitivity compared to females. Apoe^*−/−*^ mice showed minimal sex differences in atherosclerosis with a trend toward increased plaque size in females. Plaque stability markers— including collagen content, necrotic core size, and macrophage infiltration—did not differ significantly between sexes. In contrast, Ldlr^−/−^ males exhibited greater *en face* plaque burden than females, yet plaque stability remained similar across sexes.

**CONCLUSION:** Comparative analysis of two widely used murine atherosclerosis models revealed genotype-dependent sexual dimorphism. Female Apoe^−/−^ mice showed a non-significant trend toward larger plaque areas than males, whereas male Ldlr^−/−^ mice developed significantly larger *en face* atherosclerotic plaques than females. By evaluating plaque area and composition across these models, our findings underscore the importance of including both sexes in atherosclerosis studies, in accordance with guidelines from the National Institutes of Health and the American Heart Association.

---

Cardiovascular disease (CVD) remains one of the leading causes of global mortality, with significant gender disparities observed in its incidence, clinical manifestations, complications, and risk factors^1^. Women typically experience a later onset of CVD compared to men, but the prevalence in women catches up to or even surpasses that in men in older age groups, particularly in women of postmenopausal age. Additionally, women often present with a higher burden of complications and risk factors^2^. Notably, women’s coronary atherosclerotic plaques undergo an age-dependent phenotypic shift from stable fibrous lesions to vulnerable plaques, progressively reducing inter-sex differences in plaque characteristics—a transition partially attributed to post-menopausal estrogen decline^3, 4^. Understanding the mechanisms underlying these sex differences in atherosclerotic cardiovascular disease is crucial for refining diagnostic and therapeutic approaches and developing targeted treatment strategies^5^.

Current research demonstrates that sex disparities in cardiovascular disease arise from differences in sex hormones^6^, sex chromosomes^7^, lipid metabolism, and immune responses^8^, with recent studies further revealing sexually dimorphic subcellular profiles in atherosclerotic plaques^9^. However, findings across animal studies are often inconsistent, potentially due to variations in experimental models, dietary regimens, and feeding protocols. To address the variability introduced by mouse models, we employed two widely used experimental mouse models, Apoe^*−/−*^ mice and Ldlr^*−/−*^ mice, to investigate sex differences in atherosclerosis under high-fat, high-fructose and high-cholesterol diets. This approach seeks to provide more reliable and translatable insights gained from the sex disparities observed in preclinical mouse models. Moreover, these findings could provide valuable guidance for selecting appropriate mouse models in preclinical atherosclerosis research.

## MATERIALS AND METHODS

### Data availability

All supporting data are available within the article and its supplementary online files.

### Study design

In this study, two genetically modified and widely used mouse strains^10^, Apoe^*−/−*^ and Ldlr^*−/−*^ mice were utilized to investigate the effects of biological sex on the development of atherosclerosis and experiments were performed at the same time in both strains and both sexes. Mice of both genders were housed under a standard specific pathogen-free (SPF) environment maintained at approximately 22°C. The environment adhered to a 12-hour light-dark cycle. Mice were fed an atherogenic diet (AMLN diet, Dyets) (40% trans-fat, 2% cholesterol and 22% fructose)^11^ at 8 weeks old for twenty weeks to induce atherosclerosis development. At the end of the feeding period, mice were anesthetized, and tissue samples were collected for analysis. All animal procedures were conducted in the SPF Animal Husbandry Facility at the Laboratory Animal Centre, School of Life Sciences, University of Science and Technology of China (USTC). The study protocol was approved by the USTC Institutional Animal Care and Use Committee (Ethics Approval Number: USTCACUC27120124102).

### Body Composition Analysis

Prior to sampling, mice were weighed, and body composition (fat mass and lean mass) was measured using a body fat analyzer (Bruker Minispec Plus).

### Blood Cell and Serum lipid analysis

Two drops of fresh blood were collected from each mouse into anticoagulant tubes and analyzed for hematocrit composition using a fully automated blood cell analyzer (Mindray, BC-30 Vet). For serum lipid analysis, whole blood was allowed to clot at room temperature for 2 hours, followed by centrifugation at 12,000 × g for 10 minutes at 4°C. The supernatant serum was collected and stored at −80°C until further analysis. Serum levels of total cholesterol (TC), triglycerides (TG), low-density lipoprotein (LDL)-cholesterol, and high-density lipoprotein (HDL)-cholesterol were measured using a fully automated biochemical analyzer (Shenzhen Radiometer Life Technology; Chemray 240) with a commercial kit (Radiometer/Changchun Huili).

### Atherosclerosis analysis

Atherosclerotic plaque formation was assessed in the *en face* aorta and aortic sinus using Oil Red O staining^12, 13^. *En face* aortic vessels and heart tissues were fixed in 4% paraformaldehyde (PFA) for 24 hours. Perivascular fat was removed from the aorta, and the vessels were dissected to expose the inner surface. The aorta was briefly immersed in 60% isopropanol for 10 seconds, followed by staining with 0.3% Oil Red O for 10 minutes. Excess stain was washed off with 60% isopropanol, and the aorta was photographed against a white background. Plaque area was quantified using ImageJ software.

For aortic sinus analysis, fixed heart tissues were dehydrated in 20% sucrose solution for 48 hours, embedded in OCT compound, and frozen at −80°C. The aortic sinus was sectioned into 10 μm-thick slices using a cryostat (Leica, CM3050s, Germany). Plaque staining and quantification were performed as described above. Hematoxylin and eosin (H&E) staining and Masson’s trichrome staining were conducted on aortic sinus sections to analyze necrotic core area and collagen deposition, respectively. The necrotic core was defined as the cell-free area within plaques in H&E-stained sections.

### IPGTT and ITT

Glucose and insulin tolerance were assessed in mice after 13-14 weeks on the AMLN diet. For the IPGTT, mice were fasted for 16 hours and then injected intraperitoneally with a glucose solution (0.15 g/mL) (Sangon; A501991) at a dose of 1.5 g/kg body weight. Blood glucose levels were measured via tail vein sampling at 0, 15, 30, 45, 60, 90, and 120 minutes post-injection using blood glucose meter (Viva check; NB-loT).

For the ITT, mice were fasted for 4 hours and injected intraperitoneally with an insulin solution (0.05 U/mL) (Humalog; VL7516) at a dose of 0.5 U/kg body weight. Blood glucose levels were measured via tail vein sampling at the same time points as described for the IPGTT.

### Liver TC and TG analysis

Approximately 100 mg of liver tissue was weighed and placed in a microcentrifuge tube. Nine hundred microliters of isopropanol and grinding beads were added, and the tissue was homogenized using an automated grinder. The homogenate was centrifuged at 12,000 × g for 10 minutes at 4°C, and the supernatant was collected for analysis. Liver TC and TG levels were measured using the Total Cholesterol (TC) Colorimetric Assay Kit (Elabscience; E-BC-K109-M) and the Triglyceride (TG) Colorimetric Assay Kit (Elabscience; E-BC-K261-M), respectively, following the manufacturer’s instructions.

### Immunofluorescence Staining

Aortic sinus cryosections were allowed to equilibrate to room temperature for 30 minutes and then rehydrated in phosphate-buffered saline (PBS) for 5-10 minutes. Sections were permeabilized and blocked using a solution containing 5% goat serum and 0.1% Triton X-100 for 30 minutes. Primary antibodies against CD68 (BIORAD, MCA1957GA) were diluted and applied to the sections, followed by overnight incubation at 4°C. Unbound antibodies were washed away with PBS, and sections were incubated with fluorescent secondary antibodies (Alexa Fluor goat anti-rat 488, no. A48255) for 1 hour in the dark. Finally, sections were mounted using an anti-fade mounting medium containing DAPI (Beyotime Biotechnology) to counterstain nuclei. Images were acquired using a Leica Confocal Microscope (Leica, TCS SP8 X).

### Statistical Analysis

All data were tested for normality using the Shapiro-Wilk test and for homogeneity of variance using the Brown-Forsythe test. Parametric tests were applied to data meeting these assumptions, while non-parametric tests (Mann-Whitney U test) were used for data that did not meet the assumptions. Statistical analyses were performed using GraphPad Prism 9.0 software (GraphPad Software, San Diego, CA). Data are presented as mean ± standard deviation (SD), and error bars in graphs represent SD, with p<0.05 regarded as statistically significant.

## RESULTS

### Apoe^−/−^ mice showed minimal sex difference in atherosclerosis

Previous studies have reported that in regular chow-fed Apoe^*−/−*^ mice, males develop more aortic plaques than females with age, potentially due to estrogen-mediated regulation of macrophage cholesterol efflux^14^. However, conflicting evidence suggests that female Apoe^*−/−*^ mice exhibit worse vascular endothelial cell function and develop more atherosclerosis, particularly when fed a proatherogenic Western diet^15^. To further investigate whether a high-fat, high-cholesterol diet exacerbates atherosclerosis in females and to explore sex-specific metabolic and atherosclerotic characteristics, we conducted atherosclerosis modeling in both male and female Apoe^*−/−*^ mice fed an AMLN diet for 20 weeks.

Body composition analysis after 20 weeks revealed that while male mice consistently had higher body weights than females, they showed a non-significant trend toward lower fat mass (9.46 ± 3.46% vs. 11.43 ± 2.77%, P = 0.0947) (Fig. 1A). Random blood glucose levels were higher in males compared to females (8.31 ± 1.64 mmol/L vs. 5.47 ± 1.07 mmol/L, P = 0.0003) (Fig. 1A). Glucose tolerance and insulin sensitivity tests conducted at 13-14 weeks showed females demonstrated better glucose tolerance and insulin sensitivity than males (Fig. 1B). Lipid profiling after 20 weeks indicated that males had significantly higher levels of TC (14.95 ± 1.16 mmol/L vs. 11.24 ± 1.06 mmol/L, P < 0.0001), TG (2.58 ± 0.51 mmol/L vs. 1.75 ± 0.31 mmol/L, P = 0.0005), LDL-cholesterol (3.33 ± 0.26 mmol/L vs. 2.35 ± 0.32 mmol/L, P < 0.0001) and HDL-cholesterol (8.72 ± 0.88 mmol/L vs. 6.44 ± 0.65 mmol/L, P < 0.0001) compared to females (Fig. 1C). However, females showed significantly higher liver TC (57.85 ± 9.69 mmol/g vs. 45.53 ± 8.51 mmol/g, P = 0.0094) and TG (45.81 ± 6.13 mmol/g vs. 32.33 ± 6.31 mmol/g, P = 0.0002) levels compared to males (Fig. 1C). Analysis of immune cell composition in the blood revealed males had lower neutrophil percentages (43.56 ± 7.30% vs. 53.70 ± 9.12%, P = 0.0213) but higher lymphocyte percentages (49.44 ± 7.43% vs. 39.84 ± 8.36%, P = 0.0219) relative to females, with no sex difference in monocyte levels (7 ± 0.76% vs. 6.46 ± 1.16%, P = 0.3955) (Fig. 1D).

**Figure 1.**
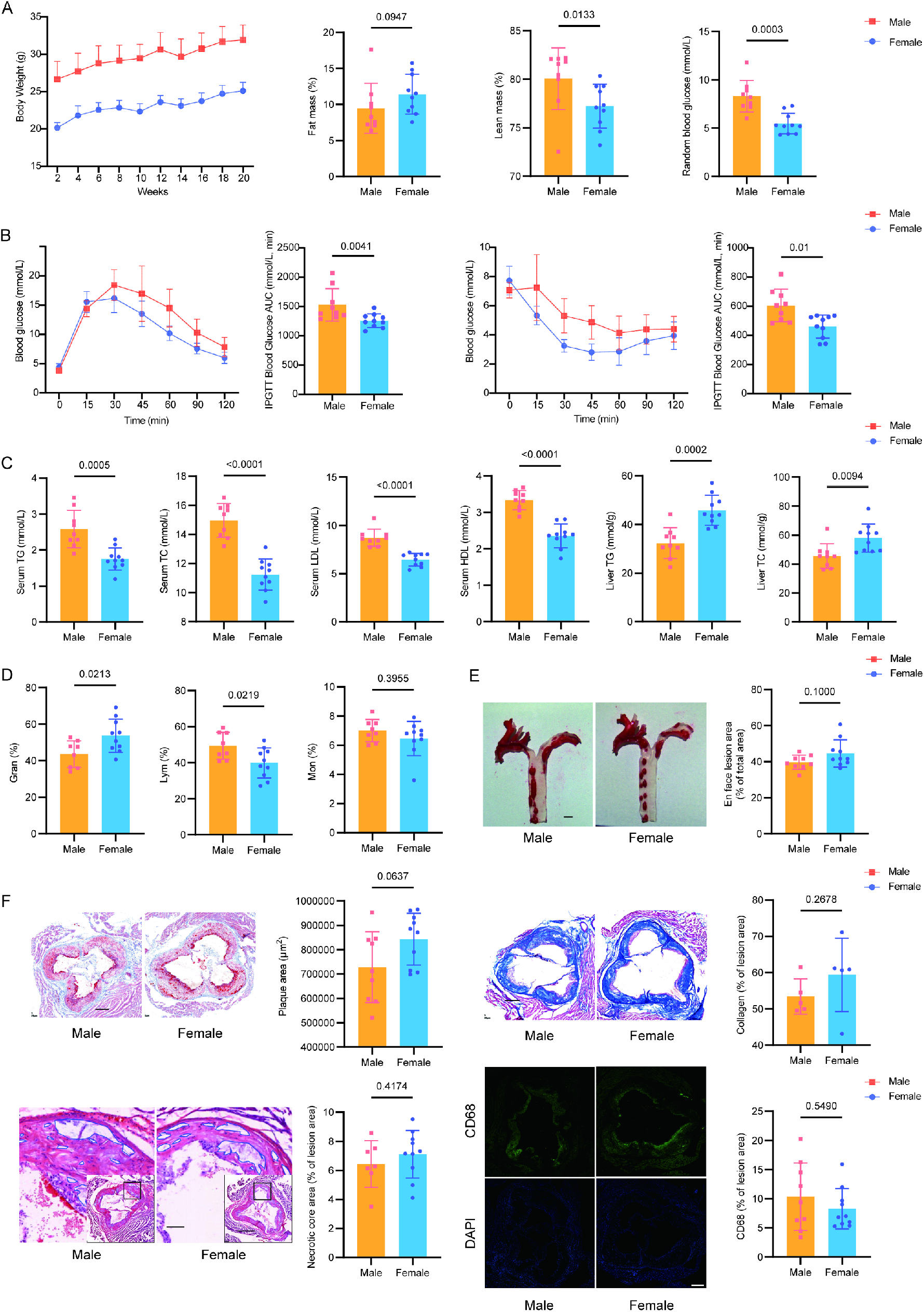
Metabolic and atherosclerotic characteristics of male and female Apoe^−/−^ mice. A. Eight-week-old male and female Apoe^−/−^ mice were fed an AMLN diet for 20 weeks to induce atherosclerosis. Fat mass, lean mass and random blood glucose were measured at endpoint (n = 9-10/group). Fat mass and lean mass were analyzed by Mann-Whitney test and random blood glucose was analyzed by unpaired t test. B. Glucose tolerance tests at week 13 (n = 9-10/group). Area under curve (AUC) comparison between sex were performed using Mann-Whitney test. Insulin tolerance tests at week 14 (n = 9-10/group). AUC comparison was analyzed by Mann-Whitney test. C. Serum lipid profiles (TC, TG, LDL, HDL) and Liver TC and TG were measured at endpoint (n = 9-10/group). unpaired t test was used in the data analyzation. D. Blood cell composition was measured at endpoint (n = 8-10/group). Gran% and Lym% were analyzed by unpaired t test. Mon% was analyzed by Mann-Whitney test. E. Aortic en face Oil Red O staining and plaque quantification (n = 9-10/group). Scale bar: 1 mm. unpaired t test was used in the data analyzation. F. Aortic sinus sections Oil Red O staining and plaque area quantification (n = 9-10/group). Scale bar: 250 μm. unpaired t test was used in the data analyzation. HE staining of aortic sinus sections and necrotic core area quantification (n = 7-10/group). Main panel scale bar: 500 μm; inset: 62.5 μm. unpaired t test was used in the data analyzation. Masson’s trichrome staining of aortic sinus sections and collagen content quantification (n = 5/group). Scale bar: 500 μm. unpaired t test was used in the data analyzation. CD68 immunofluorescence staining of aortic sinus sections and macrophage infiltration quantification (n = 9-10/group). Scale bar: 250 μm. Mann-Whitney test was used in the data analyzation. All N numbers given represent biological replicates.

Oil Red O staining of the aorta revealed that, contrary to the lipid and metabolic profiles, males didn’t have larger *en face* plaque areas than females (39.67 ± 3.96% vs. 44.58 ± 7.55%, P = 0.1000) (Fig. 1E). Similar results were observed in aortic sinus sections (728490 ± 145456 μm^2^ vs. 843727 ± 106711 μm^2^, P = 0.0637), with females exhibited a non-significant tendency toward larger plaques (Fig. 1F). Additionally, statistical analysis of necrotic core area (6.43 ± 1.61% vs. 7.1 ± 1.64%, P = 0.4174), collagen content (53.42 ± 4.88% vs. 59.4 ± 10.1%, P = 0.2678) and macrophage infiltration (10.38 ± 5.79% vs. 8.28 ± 3.45%, P = 0.5490) didn’t show any sex differences (Fig. 1F).

### The Ldlr^−/−^ male mice exhibits larger *en face* plaques than females

After 20 weeks of modeling, unlike Apoe^*−/−*^ mice, male Ldlr^*−/−*^ mice had significantly higher body fat mass than females (30.99 ± 3.02% vs. 20.16 ± 3.63%, P < 0.0001) (Fig. 2A). Although random blood glucose levels (6.6 ± 0.85 mmol/L vs. 6.76 ± 1.48 mmol/L, P = 0.7717) showed no significant differences between sexes (Fig. 2A), glucose tolerance and insulin sensitivity tests conducted at 13-14 weeks revealed that females had better glucose tolerance and insulin sensitivity than males (Fig. 2B).

**Figure 2.**
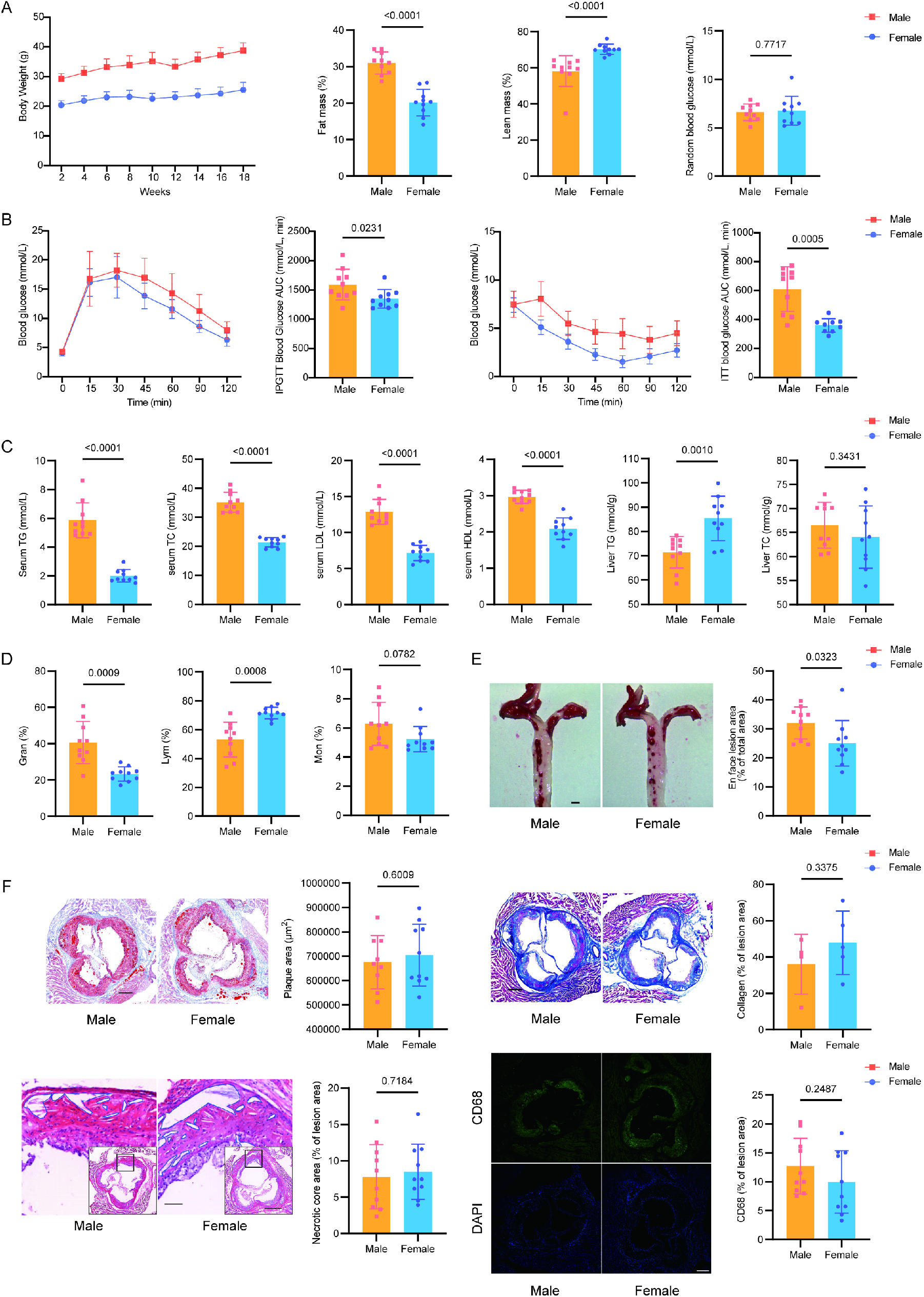
Metabolic and atherosclerotic characteristics of male and female Ldlr^−/−^ mice. A.Eight-week-old male and female Ldlr ^−/−^ mice were fed an AMLN diet for 20 weeks to induce atherosclerosis. Fat mass, lean mass and random blood glucose were measured at endpoint (n = 10/group). Fat mass and random blood glucose were analyzed by unpaired t test and lean mass was analyzed by Mann-Whitney test. B. Glucose tolerance tests at week 13 (n = 10/group). Area under curve (AUC) comparison between sex were performed using unpaired t test. Insulin tolerance tests at week 14 (n = 9-10/group). AUC comparison was analyzed by Welch’s t test. C. Serum lipid profiles (TC, TG, LDL, HDL) and Liver TC and TG were measured at endpoint (n = 10/group). TC was analyzed by Welch’s t test. TG was analyzed by Mann-Whitney test. LDL, HDL, Liver TC and Liver TG were analyzed by unpaired t test. D. Blood cell composition was measured at endpoint (n = 10/group). Gran% and Lym% were analyzed by Welch’s t test. Mon% was analyzed by Mann-Whitney test. E. Aortic en face Oil Red O staining and plaque quantification (n = 10/group). Scale bar: 1 mm. unpaired t test was used in the data analyzation. F. Aortic sinus sections Oil Red O staining and plaque area quantification (n = 9-10/group). Scale bar: 250 μm. unpaired t test was used in the data analyzation. HE staining of aortic sinus sections and necrotic core area quantification (n = 10/group). Main panel scale bar: 500 μm; inset: 62.5 μm. unpaired t test was used in the data analyzation. Masson’s trichrome staining of aortic sinus sections and collagen content quantification (n = 4-5/group). Scale bar: 500 μm. unpaired t test was used in the data analyzation. CD68 immunofluorescence staining of aortic sinus sections and macrophage infiltration quantification (n = 10/group). Scale bar: 250 μm. unpaired t test was used in the data analyzation. All N numbers given represent biological replicates.

Similar to Apoe^*−/−*^ mice, lipid profiling indicated males had significantly higher levels of TC (35.16 ± 3.4 mmol/L vs. 21.36 ± 1.63 mmol/L, P < 0.0001), TG (5.86 ± 1.22 mmol/L vs. 1.99 ± 0.44 mmol/L, P < 0.0001), LDL-cholesterol (12.88 ± 1.71 mmol/L vs. 7.16 ± 1.07 mmol/L, P < 0.0001) and HDL-cholesterol (2.96 ± 0.18 mmol/L vs. 2.08 ± 0.29 mmol/L, P < 0.0001) compared to females (Fig. 2C), and females again showed greater hepatic TG accumulation (85.41 ± 9.16 mmol/g vs. 71.46 ± 6.51 mmol/g, P = 0.001), but not TC (64.03 ± 6.46 mmol/g vs. 66.5 ± 4.74 mmol/g, P = 0.3431) (Fig. 2C). Interestingly, the leukocyte profile in Ldlr^*−/−*^ mice contrasted with Apoe^*−/−*^ mice: males had higher neutrophil (40.63 ± 11.55% vs. 23.29 ± 3.87%, P = 0.0009) and lower lymphocyte percentages (53.09 ± 12.03% vs. 71.48 ± 4.02%, P = 0.0008) compared to females, with a non-significant increase in monocytes (6.28 ± 1.47% vs. 5.23 ± 0.86%, P = 0.0782) (Fig. 2D).

Consistent with these findings, *en face* plaque area was significantly greater in male Ldlr^*−/−*^ mice (32.06 ± 5.51% vs. 25.04 ± 7.82%, P = 0.0323) (Fig. 2E), though aortic sinus plaque characteristics including area (675333 ± 109424 μm^2^ vs. 704539 ± 127329 μm^2^, P = 0.6009), collagen content (36.03 ± 16.48% vs. 47.80 ± 17.44%, P = 0.3375), necrotic core size (7.8 ± 4.42% vs. 8.47 ± 3.8%, P = 0.7184), and macrophage infiltration (12.69 ± 4.82% vs. 9.95 ± 5.4%, P = 0.2487) showed no sex-based differences (Fig. 2F).

## DISCUSSION

Sex is an important biological variable in preclinical animal-based atherosclerosis research. However, the comparative analysis of both sexes in two widely used mouse models of atherosclerosis in one study has not been conducted yet. In this study, we demonstrate that the development of atherosclerosis exhibits distinct patterns of sexual dimorphism between Apoe^*−/−*^ and Ldlr^*−/−*^ mouse models. In Apoe^*−/−*^ mice, females develop a non-significant greater atherosclerotic burden than males. In contrast, Ldlr^*−/−*^ males displayed increased *en face* plaque area compared to females. Our study carries important implications in vascular biology research, which provide unequivocal evidence in support of including both male and female mice in preclinical atherosclerosis research studies.

Our findings are consistent with a previous comparative study involving both sexes of mice, in which Apoe^*−/−*^ control mice similarly exhibited smaller atherosclerotic plaques in males compared to females, observed in both the aortic arch and sinus, with no difference in relative necrotic core area^16^. However, given that male mice exhibited less favorable lipid profiles and impaired insulin sensitivity compared to females in the present study, the metabolic parameters cannot account for the observed atheroprotection in males. This strongly suggests the involvement of other sex-specific protective mechanisms in male mice. Existing literature supports the protective role of androgens in Apoe^*−/−*^ mice. Wang et al. demonstrated that testosterone supplementation in castrated males reduced atherosclerotic plaque formation^17^, while Luo et al. identified that androgens exert cardioprotective effects by activating ADTRP transcription, thereby modulating monocyte adhesion^18^. Additionally, studies have reported that female Apoe^−/−^ mice may exhibit an enhanced immune response to oxidized low-density lipoprotein, which could contribute to accelerated atherosclerosis development^19^. Therefore, sex hormones may represent a key factor contributing to sex-based differences in atherosclerosis in Apoe^−/−^ mice. Moreover, variations in experimental outcomes across studies may arise from differences in diet, modeling duration, and the anatomical sites selected for atherosclerotic lesion assessment.

From another perspective, the AMLN diet is also used to induce metabolic dysfunction - associated steatohepatitis (MASH), which can exacerbate liver damage and influence atherosclerosis progression. Female Apoe^−/−^ mice exhibit higher hepatic levels of total cholesterol (TC) and triglycerides (TG) than males, suggesting that sex-specific lipid accumulation and associated liver injury may indirectly contribute to the progression of atherosclerosis. Furthermore, while Ldlr^−/−^ mice fed a standard high-fat diet (HFD) typically exhibit higher blood lipid levels in females than in males^6, 7, 20^, the present study revealed that Ldlr^−/−^ males fed an AMLN diet showed significantly elevated lipid levels compared to females. This may partly explain the more severe aortic plaque area observed in male Ldlr^−/−^ mice.

In summary, murine models exhibit strain-dependent sexual dimorphism in atherosclerosis development, with divergent (even opposing) sex differences observed between Apoe^*−/−*^ and Ldlr^−/−^ mice. However, both models demonstrate consistent sex-specific metabolic patterns: male mice consistently display elevated lipid profiles and reduced insulin sensitivity compared to females. Our study provides a valuable resource for the comparative analysis of glucose, lipid profile, plaque area, plaque composition in Apoe^−/−^ and Ldlr^−/−^ mice, which provides a useful resource for future atherosclerosis research.

## ARTICLE INFORMATION

## Acknowledgments

**None**

## Author contributions

Conceptualization, Suowen Xu; Investigation, Yaping Zhao; Writing – original draft, Yaping Zhao; Writing-review and editing, Li Wang, Yu Huang, Hans Strijdom, Suowen Xu, Jianping Weng and Bradford C. Berk.

